# Differential Requirements for the CENP-O Complex Reveal Parallel PLK1 Kinetochore Recruitment Pathways

**DOI:** 10.1101/2020.11.30.404327

**Authors:** Alexandra L. Nguyen, Marie Diane Fadel, Iain M. Cheeseman

## Abstract

Similar to other core biological processes, the vast majority of cell division components are essential for viability across human cell lines. However, genome-wide screens have identified a number of proteins that exhibit cell line-specific essentiality. Defining the behaviors of these proteins is critical to our understanding of complex biological processes. Here, we harness differential essentiality to reveal the contributions of the 4-subunit centromere-localized CENP-O complex, whose precise function has been difficult to define. Our results support a model in which the CENP-O complex and BUB1 act in parallel pathways to recruit a threshold level of PLK1 to mitotic kinetochores, ensuring accurate chromosome segregation. We demonstrate that targeted changes to either pathway sensitizes cells to the loss of the other component, resulting in cell-state dependent requirements. This approach also highlights the advantage of comparing phenotypes across diverse cell lines to define critical functional contributions and behaviors that could be exploited for the targeted treatment of disease.

## Introduction

A fundamental assumption for much of the research concerning core biological processes is that the conserved players that direct these processes will exhibit similar functional requirements across organisms, let alone between cell types within a given species. However, not all proteins conform to this behavior, making the identification and analysis of molecular factors with varying requirements critical to our understanding of complex cellular biology. During eukaryotic cell division, chromosomal DNA is segregated equally between daughter cells following a tightly regulated and stereotypical choreography of chromosome capture, alignment, and distribution. The key molecular players that direct chromosome segregation, including the components of the macromolecular kinetochore structure that mediates chromosome-microtubule interactions, are conserved across most eukaryotes and are essential for cellular viability (Cheeseman and Desai, 2008). Interestingly, our recent work and the results from genome-wide screens (McKinley *et al.*, 2015; Wang *et al.*, 2015; McKinley and Cheeseman, 2017; Broad, 2020) indicate that the requirement for the centromere-localized CENP-O complex varies between human cell lines. Here, we sought to exploit this cell line-specific essentiality to define the basis for these differences between cell types and the role for this complex.

The CENP-O complex is a four-subunit interdependent protein assembly, comprised of CENP-O, CENP-P, CENP-Q, and CENP-U, that localizes constitutively to centromeric DNA as part of the larger constitutive centromere-associated network (CCAN), which collectively provides the base for kinetochore assembly (Hara and Fukagawa, 2017). The viability of many human tissue culture cell lines in the absence of, the CENP-O complex is in stark contrast to other CCAN components, where perturbation results in severe mitotic defects and lethality (McKinley *et al.*, 2015). Prior work has proposed diverse functions for the CENP-O complex, including functioning as a scaffold for Polo-like kinase 1 (PLK1) recruitment to kinetochores, directly promoting kinetochore-microtubule attachments, and promoting sister chromatid cohesion (Minoshima *et al.*, 2005; Foltz *et al.*, 2006; Kang *et al.*, 2006; Kang *et al.*, 2011; Pesenti *et al.*, 2018). However, a lack of strong phenotypes observed for the loss of the CENP-O complex in the cell lines typically used for analyses of cell division, such as HeLa cells, has made defining the role of this complex difficult. Here, we harness the cell line specific requirements of the CENP-O complex to define its primary functional contribution in cellular division. By testing the cell line-specific requirements for the CENP-O complex, we find that the primary functional contribution of the CENP-O complex is in PLK1 recruitment to kinetochores. Our work reveals that PLK1 recruitment occurs through parallel pathways that are governed by BUB1 and the CENP-O complex such that changes to either pathway sensitizes cell lines to loss of the other, resulting in cell line-specific essentiality. This finding is also supported by recent work by Singh et al. (Singh *et al.*, 2020), in which the authors reconstituted the recruitment of PLK1 to *in vitro* assembled kinetochores via BUB1 and CENP-U.

Together, our work identifies the source for the differential requirement of the CENP-O complex across cell lines. Importantly, this approach also highlights the advantage of comparing differential requirements and phenotypes across diverse cell lines and cell types, particularly for defining the function of previously difficult to, characterize proteins. The investigation of cell line-specific protein essentialities will also prove valuable in pinpointing disease-specific vulnerabilities, allowing the identification of directed diagnostic and therapeutic targets.

## Results

### The CENP-O complex exhibits differential requirements in human cell lines

Despite the conservation of the 4-subunit CENP-O complex across diverse eukaryotes, our previous work found that eliminating CENP-O from human HeLa cell lines did not result in substantial defects in chromosome segregation or viability (McKinley *et al.*, 2015; McKinley and Cheeseman, 2017). Intriguingly, recent genome-wide functional screens found that the CENP-O complex is not required in most human cell lines, but also identified multiple cell lines that display a strict requirement for the CENP-O complex (Wang *et al.*, 2015; Meyers *et al.*, 2017; Broad, 2020). To define the basis for the cell line-specific requirements for the CENP-O complex, we used a Cas9-inducible gene targeting strategy (McKinley *et al.*, 2015; McKinley and Cheeseman, 2017) in multiple human cell lines. As the CENP-O complex subunits display interdependent localization (Hori *et al.*, 2008) and genome-wide functional analyses have revealed similar behaviors for each subunit (Wang *et al.*, 2015; Meyers *et al.*, 2017; Broad, 2020), for these experiments we targeted two representative CENP-O complex subunits - CENP-O and CENP-U. For our initial analysis, we compared HeLa cells, a cervical cancer cell line that our previous work found is insensitive to the loss of CENP-O, the diploid and non-transformed RPE-1 cell line, and K-562 cells, a leukemogenic cell line that exhibits proliferation defects upon gene targeting of CENP-O complex subunits based on genome-wide screens (Wang *et al.*, 2015).

We first defined the phenotypes resulting from the inducible knockout (iKO) of CENP-O or CENP-U. Due to the nature of the inducible knockout system, a subset of, Cas9 cleavage events will be repaired in a manner that retains the open reading frame resulting in a mixed population of cells. Importantly, we did not observe a difference in the proportion of cells in which the CENP-O complex was eliminated across HeLa, RPE-1, or K-562 cell line backgrounds, as determined by immunofluorescence analysis with an antibody specific for CENP-O/P (data not shown)(McKinley *et al.*, 2015), allowing us to compare behaviors between cell lines. Loss of CENP-O or CENP-U did not significantly affect chromosome alignment in HeLa or RPE-1 cells, indicating that the lack of a strong phenotype is unrelated to p53 status. In contrast, knockout of either protein resulted in dramatic mitotic defects in K-562 cells, with 39% of CENP-O iKO and 30% of CENP-U iKO cells exhibiting misaligned chromosomes, as defined by the presence of at least one off-axis chromosome, compared to 8% of control cells (Figures 1 A-B; Figure S1 A-B). Despite the presence of mis-aligned chromosomes, K-562 CENP-O and CENP-U iKO cells failed to arrest in metaphase, resulting in a proportion of cells with anaphase chromosome segregation defects, including lagging chromosomes and anaphase bridges (Figures 1 C-D; Figure S1 C-D). We note that, despite an increase in anaphase phenotypes upon CENP-O/U loss, K562 cells exhibited higher proportions of anaphase defects independent of CENP-O/U status. Together, these data highlight core differences in the sensitivity of different human cell lines to the loss of the CENP-O complex, and demonstrate that this complex contributes to chromosome alignment and segregation in specific human cell lines.

**Figure 1.**
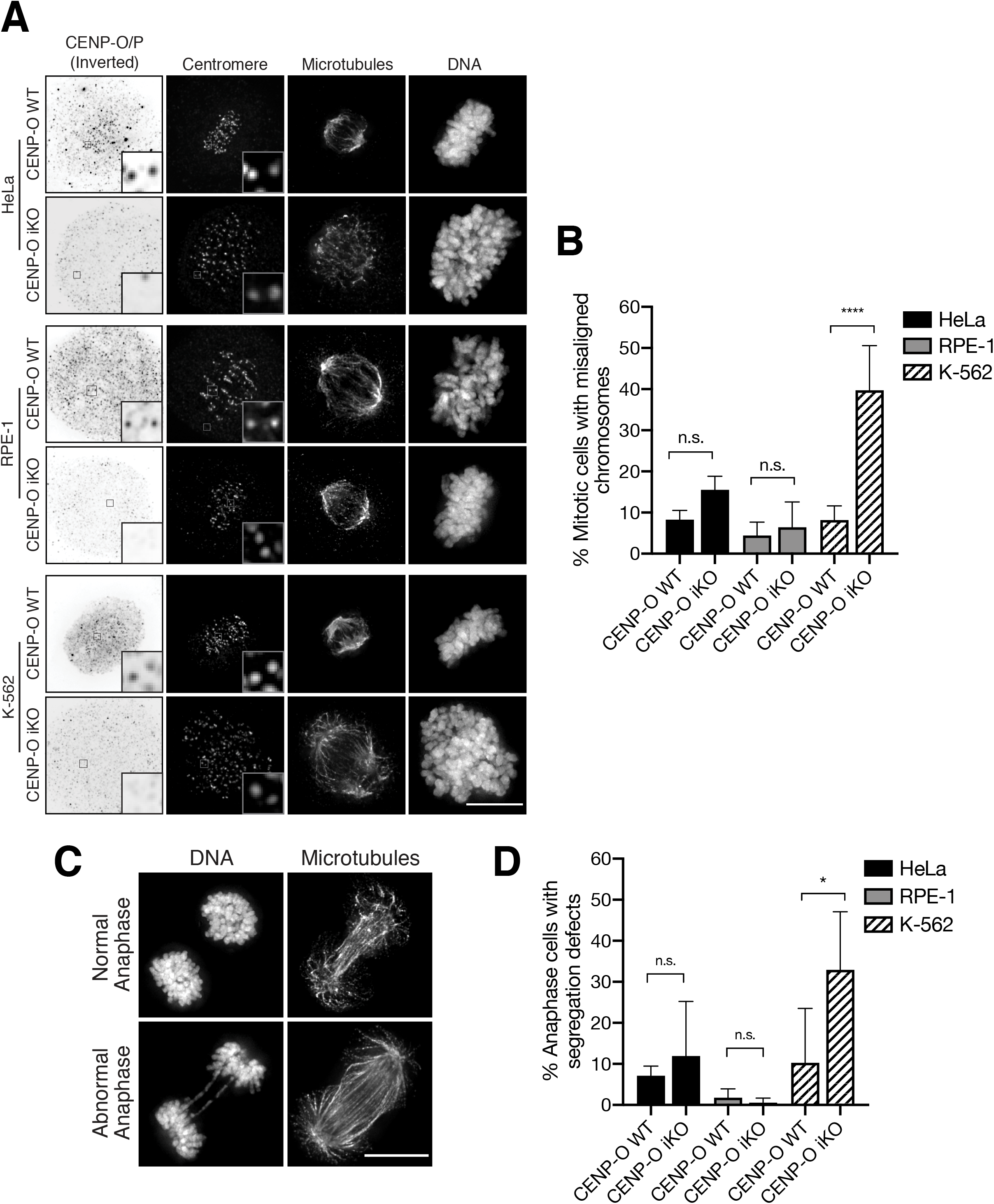
The CENP-O complex exhibits differential requirements in human cell lines. A) Representative Z-projected immunofluorescence images of metaphase cells from CENP-O inducible knockout (iKO) HeLa, RPE-1, and K-562 cell lines. Images show anti-CENP-O/P antibodies (inverted), centromeres (ACA), microtubules (DM1α), DNA (Hoechst). Boxes indicate areas of optical zoom. B) Percent mitotic cells with misaligned chromosomes after inducible knockout of CENP-O for 5 days, quantified from A. n = approximately 300 cells per condition, across 3 experimental replicates. C) Representative Z-projected immunofluorescence images of anaphase cells from CENP-O inducible knockout HeLa, RPE-1, and K-562 cell lines. Spindle (DM1α), DNA (Hoechst). D) Quantification of anaphase cells with defects including chromosome bridges and lagging chromosomes from C. Representative anaphase cells are from CENP-U control and CENP-U iKO K-562 cell lines. N = approximately 100 cells per condition across 3 experimental replicates. Error bars indicate standard deviation. One-way ANOVA was performed (* = 0.0366, **** = <0.001). Scale bars, 10 μM. See also Figure S1.

### The CENP-O complex promotes PLK1 recruitment to kinetochores

The CENP-O complex localizes constitutively to centromeres as part of the inner kinetochore Constitutive Centromere-Associated Network (CCAN) and has been proposed to perform diverse roles, including functioning as a scaffold for Polo-like kinase 1 (PLK1) recruitment to kinetochores, directly promoting kinetochore-microtubule attachments, and promoting sister chromatid cohesion (Minoshima *et al.*, 2005; Foltz *et al.*, 2006; Kang *et al.*, 2006; Elowe *et al.*, 2007; Kang *et al.*, 2011; Pesenti *et al.*, 2018). Each of these proposed functions is considered to be an essential process for all dividing cells, raising the question of why the CENP-O complex proteins would exhibit cell line-specific requirements. Prior work has focused on the functional analysis of the CENP-O complex in human cell lines where this complex is not required for viability, such as HeLa cells. Because loss of the CENP-O complex in K-562 cells results in significant increase in mitotic defects, this phenotype provides the opportunity to define the critical contributions of the CENP-O complex under conditions where it is required for cell division. The differences in CENP-O complex requirements across cell lines likely reflect underlying genetic or physiological susceptibilities that cause a given cell line to be predisposed to CENP-O complex loss. In assessing differences between cell lines, we observed a striking difference in the levels of kinetochore-localized PLK1, with reduced PLK1 levels in K-562 cells compared to HeLa and RPE-1 cells (Figure 2 A-B). The difference in PLK1 levels was specific to the kinetochore population of PLK1, as centrosomal and spindle midzone-localized PLK1 were not notably different between cell lines (Figure S2 A-B). In addition, total PLK1 protein levels were not significantly different between HeLa and K562 cells, independent of CENP-O status, suggesting that the reduced levels of kinetochore-localized PLK1 reflect differences in the recruitment of the kinase to kinetochores (Figure S2 C).

**Figure 2.**
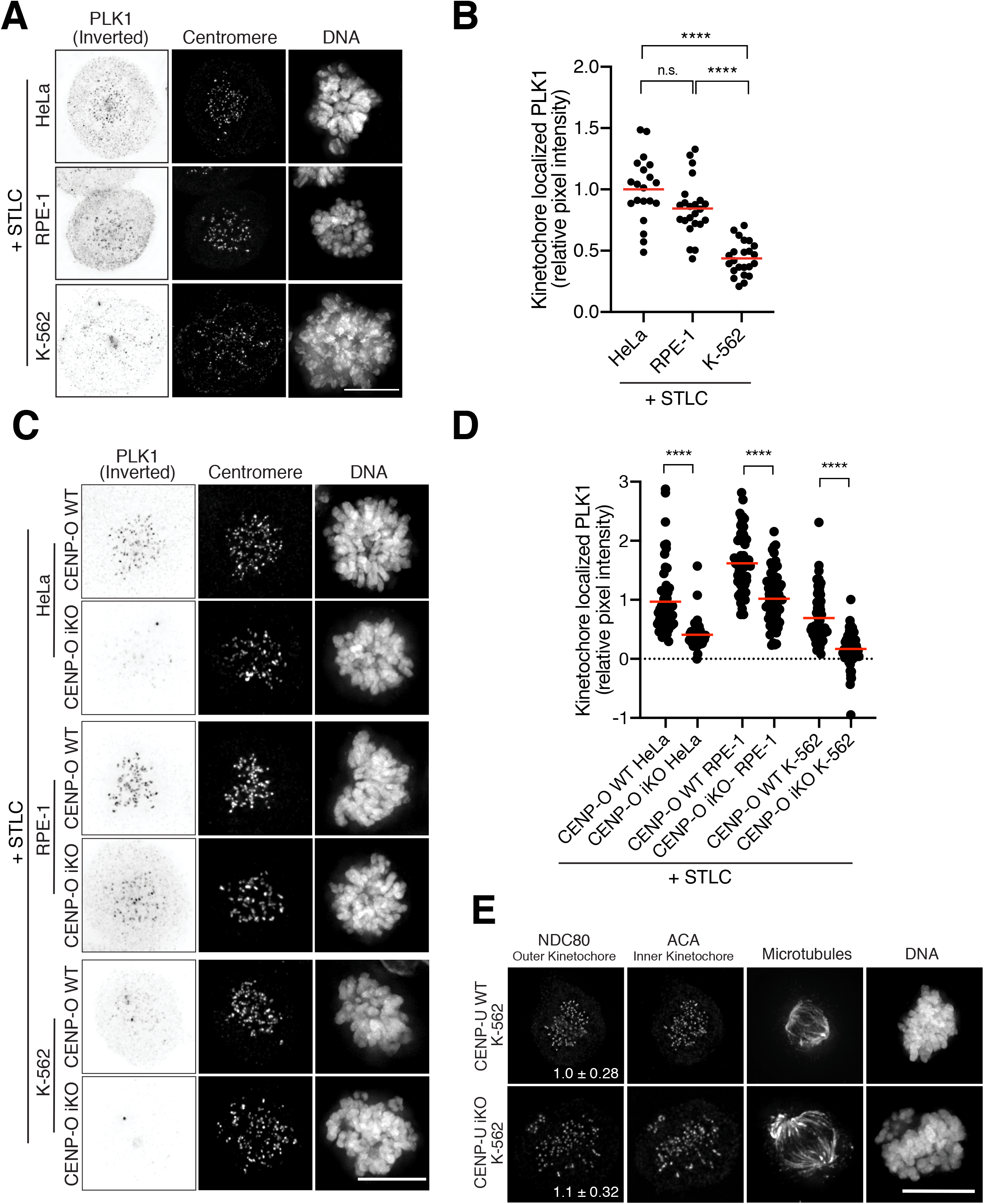
The CENP-O complex recruits PLK1 to mitotic kinetochores. A) Representative Z-projected immunofluorescence images of STLC-arrested metaphase cells from HeLa, RPE-1, and K-562 cell lines. Images show anti-PLK1 antibodies (inverted), centromeres (ACA), DNA (Hoechst). To ensure a comparison of PLK1 levels at similar stages of mitosis, cells were synchronized via incubation in the Kif11 inhibitor STLC overnight prior to fixation. B) Relative pixel intensity of kinetochore-localized PLK1, normalized to HeLa control cells, from A. Each data point represents a single cell. N = 25 cells per group across 3 experimental replicates. Red bars indicate mean. C) Representative Z-projected immunofluorescence images of STLC-arrested metaphase cells from CENP-O inducible knockout HeLa, RPE-1, and K-562 cell lines. Images show anti-PLK1 antibodies (inverted), centromeres (ACA), DNA (Hoechst). D) Relative pixel intensity of kinetochore-localized PLK1, normalized to HeLa CENP-O WT, from C. Each data point represents a single cell. Red bars indicate mean. N = approximately 60 cells per group across 2 experimental replicates. For A-D, One-way ANOVA was performed (**** = <0.0001). E) Representative Z-projected immunofluorescence images of mitotic cells from the CENP-U inducible knockout K-562 cell line after inducible knockout of CENP-U for 5 days showing NDC80, anti-centromere antibodies (ACA), microtubules (DM1α), and DNA (Hoechst). Inset ratios represent the relative pixel intensity of kinetochore-localized NDC80 ± standard deviation, normalized to control cells. N = approximately 40 cells per group across 2 experimental replicates. Student’s t-test was performed with no significant difference observed. Scale bars, 10 μM. See also Figure S2.

We next sought to test whether the varying levels of kinetochore-localized PLK1 could underlie the differential requirements for the CENP-O complex between cell lines. CENP-U binds directly to PLK1 and this binding has been proposed to promote PLK1 localization to kinetochores (Kang *et al.*, 2006; Lee *et al.*, 2008; Kang *et al.*, 2011; Park *et al.*, 2015). Based on this predicted function, we hypothesized that a threshold level of kinetochore-localized PLK1 is required for accurate chromosome segregation, and that cell lines with reduced PLK1 would be sensitized to CENP-O/U depletion. Consistent with this hypothesis, we found that eliminating CENP-O or CENP-U resulted in a significant reduction in kinetochore-localized PLK1 in all the cell lines tested when compared to controls from the corresponding cell line (Figures 2 C-D and S2 D-E). Despite all cell lines exhibiting a reduction in PLK1 levels upon CENP-O/U depletion, the level of PLK1 at kinetochores was significantly lower in K562 CENP-O and CENP-U iKO cells when compared to both HeLa CENP-O/U iKO or RPE-1 CENP-O/U iKO cells. Although Plk1 levels were variable across experiments, we observed a consistent trend in which a large proportion of K562 cells exhibited substantially lower PLK1 levels than either HeLa or RPE-1 cells, with a greater spread observed across the population. In contrast to PLK1 localization, the localization of the outer kinetochore component NDC80 was not affected by CENP-U loss (Figure 2E), indicating that the mitotic defects observed upon the loss of the CENP-O complex in K-562 cells are not the result of kinetochore assembly defects. These results support a model in which the CENP-O complex promotes the recruitment of PLK1 to mitotic kinetochores.

Although eliminating CENP-O or CENP-U results in a reduction in kinetochore-localized PLK1, PLK1 localization is not lost completely suggesting that additional kinetochore-localized PLK1 binding partners contribute to its recruitment (Figure 2C-D; Figure S2 D-E) (Kang *et al.*, 2006; Lee *et al.*, 2008; Kang *et al.*, 2011). In addition to CENP-U (Kang *et al.*, 2006; Lee *et al.*, 2008), multiple kinetochore-localized proteins have been proposed to serve as PLK1 binding factors, including BUB1 and INCENP (Goto *et al.*, 2006; Kang *et al.*, 2006; Qi *et al.*, 2006). Therefore, we next investigated whether these alternate PLK1 recruitment pathways were altered in K-562 cells, creating a synthetic lethal-like relationship for the CENP-O complex. Interestingly, we found that BUB1 levels, but not INCENP levels, were significantly lower at kinetochores in K-562 cells when compared to HeLa or RPE-1 (Figure 3 A-C). Consistently, total BUB1 protein levels also appeared mildly reduced in K-562 cells, as compared to HeLa (Figure S 2C), suggesting that the differential expression of kinetochore-localized binding partners for PLK1 could underlie the cell-type specific requirement for the CENP-O complex.

**Figure 3.**
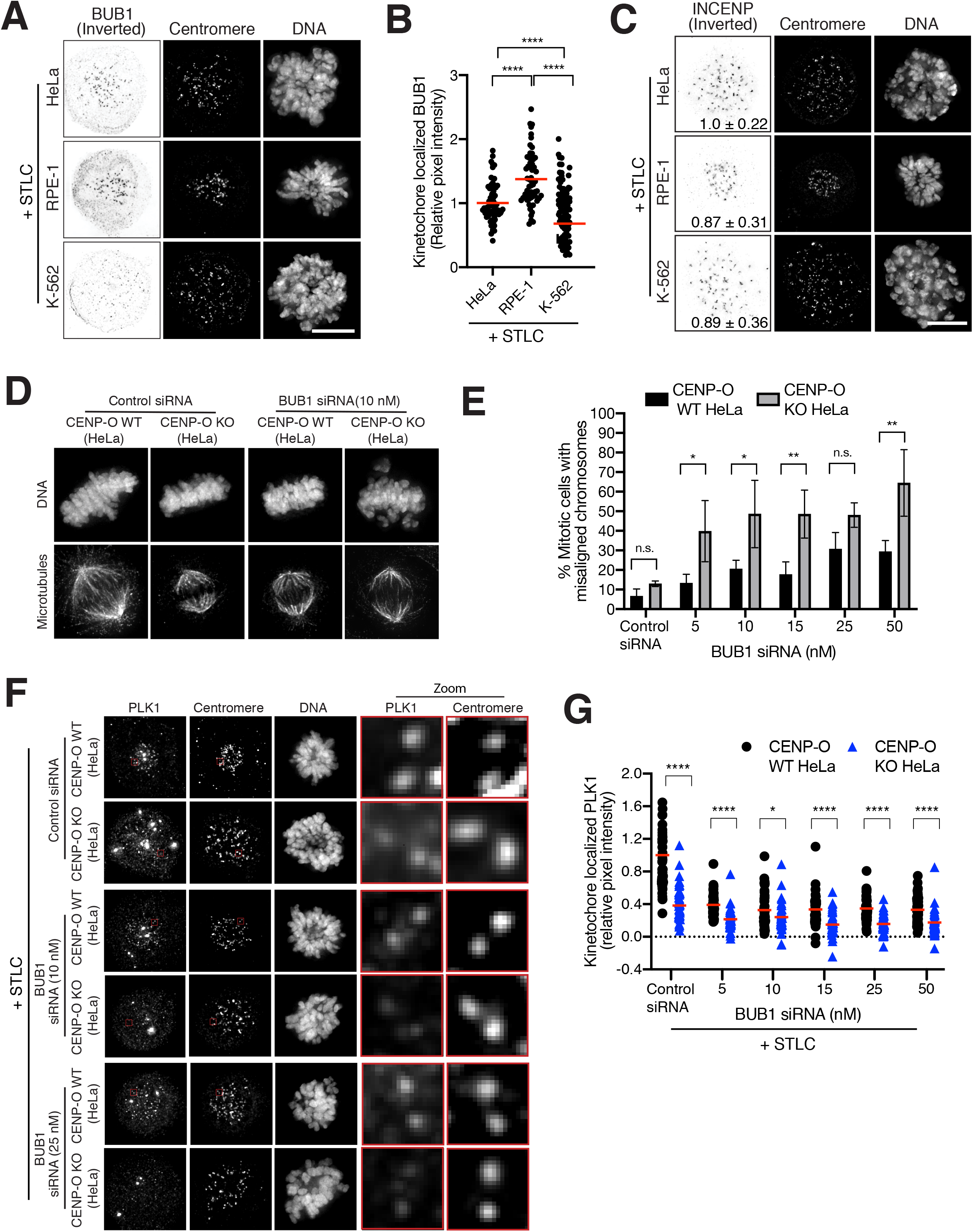
Reducing BUB1 expression sensitizes cells to the loss of the CENP-O complex. A) Representative Z-projected immunofluorescence images of STLC-arrested metaphase cells from HeLa, RPE-1, and K-562 cell lines showing anti-BUB1 antibodies (inverted), centromeres (ACA), and DNA (Hoechst). To ensure a comparison of protein levels at similar stages of mitosis, cells were synchronized via incubation in the Kif11 inhibitor STLC overnight prior to fixation. B) Relative pixel intensity of kinetochore-localized BUB1, normalized to HeLa, from A. Each data point represents a single cell. Red bars indicate the mean. N = approximately 60 cells per group across 3 experimental replicates. One-way ANOVA was performed (**** < 0.0001). C) Representative Z-projected immunofluorescence images of STLC-arrested metaphase cells from HeLa, RPE-1, and K-562 cell lines showing anti-INCENP (inverted), centromeres (ACA), and DNA (Hoechst). Inset ratios represent the relative pixel intensity of kinetochore-localized INCENP ± standard deviation, normalized to HeLa. N = approximately 30 cells per group across 2 experimental replicates. One-way ANOVA was performed with no significant difference observed. D) Z-projected immunofluorescence images of metaphase cells of the indicated cell lines incubated in the presence of control siRNA or 10 nM BUB1 siRNA showing microtubules (DM1α) and DNA (Hoechst). HeLa CENP-O WT and stable CENP-O knockout (KO) cells were incubated in the presence of the indicated concentrations BUB1 siRNA or non-targeting control for 48 hours prior to analysis. E) Percent mitotic cells with misaligned chromosomes from D. Error bars indicate standard deviation. N = approximately 300 cells per condition/per group, across 3 experimental replicates. Two-way ANOVA was performed ((5nM)* = 0.02, (10nM)* = 0.01, (15nM)** = 0.006, (50 nM)** = 0.001). F) Representative Z-projected immunofluorescence images of STLC-arrested metaphase cells of the indicated cell lines incubated in the presence of control siRNA and 10 nM BUB1 siRNA showing anti-PLK1 antibodies, centromeres (ACA), and DNA (Hoechst). To ensure a comparison of PLK1 levels at similar stages of mitosis, cells were synchronized via incubation in the Kif11 inhibitor STLC overnight prior to fixation. Boxes indicate areas of optical zoom. G) Relative pixel intensity of kinetochore-localized PLK1 from F, normalized to control siRNA CENP-O WT HeLa. N = approximately 50 cells per group, across 2 experimental replicates. Red bars indicate the mean. Statistics represent T-test comparing control and CENP-O knockout PLK1 measures per concentration siRNA (* = 0.03, **** = <0.001). Scale bars, 10 μM. See also Figure S3.

### The CENP-O complex and BUB1 collaborate to recruit PLK1 to mitotic kinetochores

Based on the data described above, we hypothesized that BUB1 and the CENP-O complex act in parallel to recruit PLK1 to mitotic kinetochores such that cell lines with reduced kinetochore-localized BUB1 would have an increased requirement for the CENP-O complex. To test this model, we sought to sensitize cell lines in which the CENP-O complex is otherwise dispensable and rescue cell lines that normally require the CENP-O complex. To this end, we first generated varying levels of BUB1 using partial RNAi-based depletion in HeLa cells, a cell line that is not normally sensitive to CENP-O loss. Due to the insensitivity of K-562 cells to siRNA treatment via standard transfection techniques, independent of the target sequence, we were unable to conduct similar experiments in this cell background. Strikingly, CENP-O knockout HeLa cells were hypersensitive to the reduction in BUB1 levels, with concentrations as low as 5 nM BUB1 siRNA resulting in a significant increase in chromosome misalignment (Figure 3D-E). This phenotype is in stark contrast to control cells, in which only a modest increase in chromosome alignment was observed at concentrations below 25 nM BUB1 siRNA. The increase in mitotic defects observed upon BUB1 depletion correlated with a dose-dependent reduction in kinetochore-localized PLK1 in both control and CENP-O KO HeLa cells, consistent with a role for BUB1 in promoting PLK1 kinetochore localization (Figure 3F-G). The sensitivity of CENP-O knockout HeLa cells to PLK1 loss was specific to BUB1 perturbation, as knockdown of INCENP did not result in a significant increase in chromosome mis-alignment in CENP-O knockout HeLa cells when compared to controls (Figure S3 A-B). Notably, at the concentrations of INCENP siRNA employed in this study, no significant difference in kinetochore-localized PLK1 was observed between control and siRNA knockdown cells (Figure S3 C-D). Furthermore, these effects were not the result of a deficient spindle assembly checkpoint, because partial depletion of MAD2 resulted in comparable defects in both control and CENP-O knockout HeLa cells (Figure S3E-F). Together, these data support a model in which BUB1 and the CENP-O complex cooperate to recruit PLK1 to mitotic kinetochores such that altering either pathways, creates an increased reliance on the other pathway to ensure sufficient PLK1 localization to kinetochores.

Next, we sought to suppress the requirement for the CENP-O complex in cell lines in which it is normally required for chromosome segregation. If the requirement for the CENP-O complex in K-562 cells reflects reduced BUB1 levels, we reasoned that increased expression of BUB1 should be sufficient to suppress the chromosome segregation defects observed upon CENP-O complex loss in K-562 cells. Indeed, stable expression of GFP-BUB1 was sufficient to rescue the mitotic defects observed upon inducible knockout of CENP-U in K-562 cells, with levels of chromosome misalignment comparable to control cells (Figure 4A-B). Similarly, expression of GFP-BUB1 resulted in a significant increase in PLK1 kinetochore localization in CENP-U inducible KO K-562 cells, comparable to those observed in control cells (Figure 4C-D). Together, these data support a model in which BUB1 and the CENP-O complex act in parallel to ensure that a threshold level of PLK1 is localized to mitotic kinetochores.

**Figure 4.**
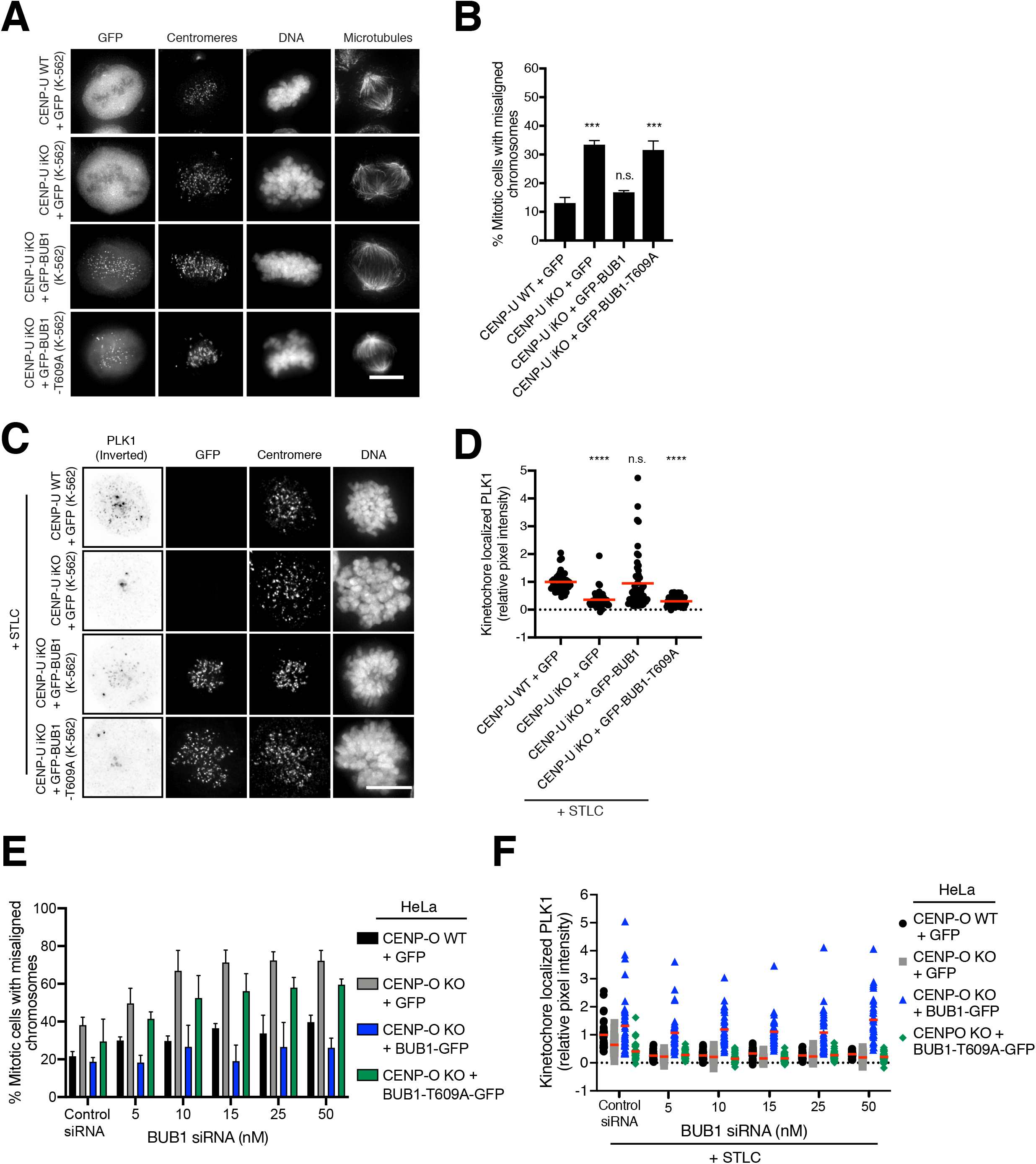
Ectopic BUB1 expression can suppress the cell-line specific requirement for the CENP-O complex. A) Representative Z-projected immunofluorescence images of metaphase cells from CENP-U inducible knockout K-562 cell lines expressing the indicated constructs showing GFP, centromeres (ACA), DNA (Hoechst), and microtubules (DM1α). B) Percent mitotic cells with misaligned chromosomes after inducible knockout of CENP-U for 5 days, from A. N = approximately 250 cells per condition, across 3 experimental replicates. Error bars indicate standard deviation. One-way ANOVA performed ((CENP-U iKO + GFP)***=0.0007, (CENP-U iKO + GFP-BUB1-T609A)*** = 0.0002). C) Representative Z-projected immunofluorescence images of STLC-arrested metaphase cells from CENP-U inducible knockout K-562 cell lines expressing the indicated constructs showing anti-PLK1 antibodies (inverted), GFP, DNA (Hoechst), and microtubules (DM1α). To ensure a comparison of PLK1 levels at similar stages of mitosis, cells were synchronized via incubation in the Kif11 inhibitor STLC overnight prior to fixation. D) Relative pixel intensity of kinetochore-localized PLK1, normalized to CENP-U WT + GFP control, from C. Each data point represents a single cell. Red bars indicate mean. N = Approx. 40 cells per group across 2 experimental replicates. One-way ANOVA was performed (**** < 0.0001). E-F) HeLa control and stable CENP-O knockout (KO) cells expressing the indicated constructs were incubated in the presence of the indicated concentrations BUB1 siRNA or non-targeting control for 48 hours prior to analysis. E) Percent mitotic cells with misaligned chromosomes from the indicated cell lines after 48 hours BUB1 siRNA knockdown. Error bars indicate standard deviation. N = approximately 300 cells per condition/per group, across 3 experimental replicates. See Table 1 for summary of corresponding statistical analysis. F) Relative pixel intensity of kinetochore-localized PLK1 in the indicated cell lines after 48 hours BUB1 siRNA knockdown followed by overnight incubation in STLC, normalized to control siRNA CENP-O WT + GFP HeLa. N = approximately 50 cells per group, across 2 experimental replicates. See Table 2 for summary of corresponding statistical analysis. Red bars indicate mean. Scale bars, 10 μM.

### Defects in PLK1-kinetochore recruitment underlie the chromosome segregation defects in CENP-O knockout cells

BUB1 plays diverse roles in the control of cell division in addition to its ability to recruit PLK1 to kinetochores (Marchetti and Venkatachalam, 2010; Combes *et al.*, 2017). To determine whether insufficient recruitment of PLK1 to mitotic kinetochores underlies the defects observed in CENP-O-complex-depleted cells, we generated a mutation in BUB1 (T609A) that prevents PLK1 binding without interfering with other known activities (Qi *et al.*, 2006). In K-562 cells, which display a requirement for the CENP-O complex, expression of GFP-BUB1 rescued the mitotic defects and the corresponding reduction in kinetochore-localized PLK1 observed upon depletion of CENP-U (Figure 4A-D). In contrast, the PLK1 binding-deficient BUB1 mutant (T609A) failed to rescue these defects (Figure 4A-D), highlighting the functional requirement for PLK1 recruitment. Similarly, we tested whether expression of RNAi-resistant versions of GFP-BUB1 or GFP-BUB1-T609A were able to rescue the defects caused by partial depletion of BUB1 by RNAi in HeLa CENP-O knockout cells. Expression of BUB1-GFP, but not BUB1-T609A-GFP, was sufficient to rescue the chromosome alignment defects and the reduced levels of kinetochore-localized PLK1 observed in CENP-O HeLa KO cells treated with BUB1 siRNAs (Figure 4E-F, Table 1,Table 2). Taken together, these data suggest that BUB1 and the CENP-O complex act in parallel pathways to recruit a threshold level of PLK1 to kinetochores. The presence of either pathway is sufficient to promote PLK1 function at kinetochores, but perturbations to one of these proteins generates a synthetic lethal requirement for the other to ensure a threshold level of PLK1 is maintained. Failure to recruit a minimum level of PLK1 results in severe chromosome segregation defects.

**Table 1.** Pair-wise comparisons of percent chromosome misalignment in the indicated cell lines transfected with the indicated concentrations siRNA. Data shown in Figure 4E. Dunnet’s multiple comparison test used for statistical analysis of groups.

| Group 1 + siRNA [nM] | Group 2 + siRNA [nM] | Significance summary | Adjusted P-Value |
| --- | --- | --- | --- |
| CENP-O WT+ GFP [0 nM] | CENP-O KO + GFP [0 nM] | ns | 0.0986 |
| CENP-O WT+ GFP [0 nM] | CENP-O KO + BUB1-GFP [0 nM] | ns | 0.9994 |
| CENP-O WT+ GFP [0 nM] | CENPO KO + BUB1-T609A-GFP [0 nM] | ns | 0.9093 |
| CENP-O WT+ GFP [0 nM] | CENP-O WT + GFP [5 nM] | ns | 0.695 |
| CENP-O WT+ GFP [0 nM] | CENP-O KO + BUB1-GFP [5 nM] | *** | 0.0005 |
| CENP-O WT+ GFP [0 nM] | CENPO KO + BUB1-T609A-GFP [5 nM] | ns | 0.9993 |
| CENP-O WT+ GFP [0 nM] | CENP-O KO + BUB1-GFP [5 nM] | * | 0.0249 |
| CENP-O WT+ GFP [0 nM] | CENP-O WT+ GFP [10 nM] | ns | 0.7282 |
| CENP-O WT+ GFP [0 nM] | CENP-O KO + GFP [10 nM] | **** | <0.0001 |
| CENP-O WT+ GFP [0 nM] | CENP-O KO + BUB1-GFP [10 nM] | ns | 0.9989 |
| CENP-O WT+ GFP [0 nM] | CENPO KO + BUB1-T609A-GFP [10 nM] | *** | 0.0001 |
| CENP-O WT+ GFP [0 nM] | CENP-O WT+ GFP [15 nM] | ns | 0.0699 |
| CENP-O WT+ GFP [0 nM] | CENP-O KO + GFP [15 nM] | **** | <0.0001 |
| CENP-O WT+ GFP [0 nM] | CENP-O KO + BUB1-GFP [15 nM] | ns | 0.9995 |
| CENP-O WT+ GFP [0 nM] | CENPO KO + BUB1-T609A-GFP [15 nM] | *** | 0.0002 |
| CENP-O WT+ GFP [0 nM] | CENP-O WT+ GFP [25 nM] | ns | 0.2232 |
| CENP-O WT+ GFP [0 nM] | CENP-O KO + GFP [25 nM] | **** | <0.0001 |
| CENP-O WT+ GFP [0 nM] | CENP-O KO + BUB1-GFP [25 nM] | ns | 0.9989 |
| CENP-O WT+ GFP [0 nM] | CENPO KO + BUB1-T609A-GFP [25 nM] | **** | <0.0001 |
| CENP-O WT+ GFP [0 nM] | CENP-O WT+ GFP [50 nM] | * | 0.0245 |
| CENP-O WT+ GFP [0 nM] | CENP-O KO+ GFP [50 nM] | **** | <0.0001 |
| CENP-O WT+ GFP [0 nM] | CENP-O KO + BUB1-GFP [50 nM] | ns | 0.999 |
| CENP-O WT+ GFP [0 nM] | CENPO KO + BUB1-T609A-GFP [50 nM] | **** | <0.0001 |

Dunnet's multiple comparisons test Figure 4E

**Table 2.** Pair-wise comparisons of kinetochore-localized PLK in the indicated cell lines transfected with the indicated siRNA concentrations. Values are indicated relative to control cells. Data shown in Figure 4F. Tukey’s multiple comparison test used for statistical analysis of groups.

Kinetochore-localized PLK1 (Relative pixel intensity)
| Group 1 + siRNA [nM] | Group 2 + siRNA [nM] | Significance summary | Adjusted P-Value |
| --- | --- | --- | --- |
| CENP-O WT+ GFP [0 nM] | CENP-O KO + GFP [0 nM] | * | 0.0271 |
| CENP-O WT+ GFP [0 nM] | CENP-O KO + BUB1-GFP [0 nM] | ns | 0.0796 |
| CENP-O WT+ GFP [0 nM] | CENPO KO + BUB1-T609A-GFP [0 nM] | **** | <0.0001 |
| CENP-O WT+ GFP [0 nM] | CENP-O WT + GFP [5 nM] | **** | <0.0001 |
| CENP-O WT+ GFP [0 nM] | CENP-O KO + BUB1-GFP [5 nM] | **** | <0.0001 |
| CENP-O WT+ GFP [0 nM] | CENPO KO + BUB1-T609A-GFP [5 nM] | **** | <0.0001 |
| CENP-O WT+ GFP [0 nM] | CENP-O KO + BUB1-GFP [5 nM] | ns | >0.9999 |
| CENP-O WT+ GFP [0 nM] | CENP-O WT+ GFP [10 nM] | **** | <0.0001 |
| CENP-O WT+ GFP [0 nM] | CENP-O KO + GFP [10 nM] | **** | <0.0001 |
| CENP-O WT+ GFP [0 nM] | CENP-O KO + BUB1-GFP [10 nM] | ns | 0.949 |
| CENP-O WT+ GFP [0 nM] | CENPO KO + BUB1-T609A-GFP [10 nM] | **** | <0.0001 |
| CENP-O WT+ GFP [0 nM] | CENP-O WT+ GFP [15 nM] | **** | <0.0001 |
| CENP-O WT+ GFP [0 nM] | CENP-O KO + GFP [15 nM] | **** | <0.0001 |
| CENP-O WT+ GFP [0 nM] | CENP-O KO + BUB1-GFP [15 nM] | ns | >0.9999 |
| CENP-O WT+ GFP [0 nM] | CENPO KO + BUB1-T609A-GFP [15 nM] | **** | <0.0001 |
| CENP-O WT+ GFP [0 nM] | CENP-O WT+ GFP [25 nM] | **** | <0.0001 |
| CENP-O WT+ GFP [0 nM] | CENP-O KO + GFP [25 nM] | **** | <0.0001 |
| CENP-O WT+ GFP [0 nM] | CENP-O KO + BUB1-GFP [25 nM] | ns | >0.9999 |
| CENP-O WT+ GFP [0 nM] | CENPO KO + BUB1-T609A-GFP [25 nM] | **** | <0.0001 |
| CENP-O WT+ GFP [0 nM] | CENP-O WT+ GFP [50 nM] | **** | <0.0001 |
| CENP-O WT+ GFP [0 nM] | CENP-O KO+ GFP [50 nM] | **** | <0.0001 |
| CENP-O WT+ GFP [0 nM] | CENP-O KO + BUB1-GFP [50 nM] | **** | <0.0001 |
| CENP-O WT+ GFP [0 nM] | CENPO KO + BUB1-T609A-GFP [50 nM] | **** | <0.0001 |

Tukey's multiple comparisons test Figure 4F

## Discussion

### Parallel PLK1 kinetochore recruitment pathways underlie differential CENP-O complex requirements

The functional contributions of the CENP-O complex to cell division have been difficult to define, in part due to the insensitivity of many cell lines to its loss. Here, using a combination of cell biological and genetic approaches, we find that a primary functional contribution for the CENP-O complex in human cells is to recruit PLK1 to mitotic kinetochores. The role of PLK1 at mitotic kinetochores has been of great interest (Lera *et al.*, 2019). However, as PLK1 maintains multiple distinct localizations, and plays diverse roles during mitosis, including in centrosome function, cytokinesis, spindle orientation, and other tasks (Colicino and Hehnly, 2018), strategies that globally inhibit PLK1 are unable to reveal the precise functions of PLK1 at kinetochores. Because the specific mechanisms of PLK1 recruitment to kinetochores have remained elusive, so have its kinetochore contributions. Importantly, our work demonstrates that PLK1 kinetochore localization is dependent upon parallel BUB1 and CENP-U-based recruitment pathways that together ensure a threshold level of PLK1 localization to kinetochores, promoting accurate chromosome alignment and segregation. Perturbations to either of these pathways result in a sensitized requirement for the remaining PLK1 binding partner. This is consistent with previous work that found that even subtle reductions in PLK1 activity can severely impact chromosome congression, indicating distinct activity thresholds are required for proper kinase function (Al-Bassam *et al.*, 2012). Additionally, recent work, from Singh et al further supports a model in which BUB1 and CENP-U serve as the primary recruitment pathways for PLK1 to mitotic kinetochores (Singh *et al.*, 2020).

This work also highlights an important role for PLK1 recruitment to kinetochores in mitotic chromosome alignment and segregation. Specifically, in situations with reduced PLK1 localization to kinetochores, chromosomes are unable to align at the metaphase plate and display defective segregation in anaphase (Figures 1A-D, 2A-D, S1A-D, and S2D-E). These results are consistent with prior work in which the specific inactivation of inner-kinetochore localized PLK1 disrupted chromosome alignment and segregation (Lera *et al.*, 2016), and provide evidence for CENP-U being the primary inner kinetochore PLK1 binding partner. Importantly, we find that the differential requirement exhibited by the CENP-O complex in K-562 cells, when compared to HeLa and RPE-1 cell lines, reflects differences in the BUB1-PLK1 recruitment pathway. The basis for the altered BUB1 levels remains unknown and will be an important topic for future work.

Core cellular processes such as cell division is typically considered to display similar mechanisms and requirements across cell lines. In contrast, these findings support the existence of diverse cell division behaviors and requirements across human cell lines. This work also highlights the advantage of comparing differential requirements across cell lines or cell types to determine the underlying basis for core cellular processes, including the functions of proteins that historically have been difficult to characterize. Cell line-specific susceptibilities can also identify vulnerabilities that could be exploited to screen for and develop treatment strategies for difficult to manage diseases. Such strategies could be especially beneficial in the treatment of acute myeloid leukemias (AML), cancers, that have been notoriously difficult to treat. The CENP-O complex exhibits a unique requirement in multiple AML cell lines (Wang *et al.*, 2015; Tzelepis *et al.*, 2016; Broad, 2020) and the use of kinase inhibitors that target BUB1 or PLK1 could provide be an effective strategy for treatment of these cancers while limiting the effect on normal, non-cancerous cell lines.

## Materials and Methods

### Contact for Reagent and Resource Sharing

Further information and requests for resources and reagents should be directed to and will be fulfilled by the Lead Contact, Iain Cheeseman.

### Cell Culture

The inducible Cas9 hTERT RPE-1(cTT33.1), HeLa (cTT20.11), and K-562 (cKC363) cell lines were generated by transposition as described previously (McKinley *et al.*, 2015; McKinley and Cheeseman, 2017) and are neomycin resistant. Cell lines were tested monthly for mycoplasma contamination. Inducible knockouts for CENP-O and CENP-U in HeLa (CENP-O:cKM160, CENP-U:cALN42), RPE-1 (CENP-O: cALN64, CENP-U:cALN66), and K-562 (CENP-O: cALN4, CENP-U: cALN153) cell lines were created by cloning and introducing pLenti-sqRNA (puromycin resistant) (McKinley *et al.*, 2015) into the inducible Cas9 cell lines by lentiviral transduction (Wang *et al.*, 2015) using sgRNAs targeting CENP-O (CACCGTTTACGGGATCTGCTCACT) or CENP-U (CACCGAGACTTACTGATGCTCTAGG) (McKinley *et al.*, 2015). Cells were then selected with 0.35 mg/mL (HeLa), 3 mg/mL (RPE-1), or 3 mg/mL (K-562) puromycin for 14 days. The HeLa CENP-O stable knockout cell line (cKM212) was described previously (McKinley *et al.*, 2015).

Clonal cell lines expressing GFP^LAP^ fusions for BUB1, and BUB1-T609A were generated using retroviral infection in HeLa and K-562 cells as described previously (Cheeseman and Desai, 2005). The BUB1 and BUB1-T609A templates are resistant to, siRNA targeted by mutation of the utilized siRNA target sequence (CTG TAC ATT GCC TGG GCG GGG to CTC TAT ATC GCT TGG GCC GGA). HeLa CENP-O knockout (cKM212) and control (cTT20.11), or K-562 CENP-U inducible knockout (cALN153) and control (cKC363) cell lines were transfected with retrovirus carrying the transgenes (pIC242: GFP, pALN24: GFP-BUB1, pALN25: GFP-BUB1-T609A) and selected with 2 mg/ml (HeLa) or 8 mg/ml (K-562) Blasticidin (Life Technologies) (Cheeseman and Desai, 2005).

HeLa and RPE-1 cell lines were cultured in Dulbecco’s Modified Eagle Medium (DMEM) supplemented with 10% tetracycline-free fetal bovine serum (FBS), 100 units/mL penicillin, 100 units/mL streptomycin, and 2 mM L-glutamine (complete media) at 37°C with 5% CO2. K562 cell lines were cultured in Roswell Park Memorial Institute (RPMI) 1640 medium supplemented with 10% tetracycline-free fetal bovine serum (FBS), 100 units/mL penicillin, 100 units/mL streptomycin, and 2 mM L-glutamine at 37°C with 5% CO2. For knockout experiments, HeLa cells were plated on polylysine coated coverslips, or uncoated coverslips for hTERT-RPE-1 cell lines, and 1 μg/mL doxycycline hyclate (Sigma) was added to cells at 24 hr. intervals for 3 days, with fixation on the fifth day. K-562 cell lines were cultured in the absence of coverslips, with doxycycline added as described above. On the fifth day, K-562 cell lines were adhered to polylysine coated coverslips via centrifugation at 2250 RPM for 30 mins at 37°C, followed by incubation at 37°C with 5% CO2 for 1 hour prior to fixation.

### siRNAs and drug treatment

siRNAs against BUB1 (GAGUGAUCACGAUUUCUAUUU), INCENP (UGACACGGAGAUUGCCAACUU), and MAD2 (UACGGACUCACCUUGCUUGUU) and a non-targeting control were obtained from Dharmacon. RNAi experiments were conducted using Lipofectamine RNAi MAX and reduced serum OptiMEM (Life Technologies). Media was replaced with complete media 24 hours after siRNA addition. Cells were assayed 48 hours after transfection. To synchronize cells in mitosis, S-trityl-L-cysteine (STLC) was added cells at 10 μM overnight.

### Immunofluorescence, microscopy, and Western Blotting

Cells on coverslips were fixed in 0.5% Triton-X-100 + 4% formaldehyde for 10 minutes at room temperature. Coverslips were then washed with PBS containing 0.1% Triton X-100, then blocked with AbDil (3% BSA, 1 X TBS, 0.1% Triton X-100, 0.1% Na Azide) for 30 min. Immunostaining was performed by incubating coverslips in primary antibody diluted in AbDil for 1 h at room temperature followed by 3 consecutive washes in PBS containing 0.1% Triton X-100. After washing, secondary antibodies were diluted 1:300 in Abdil and the sample was incubated for 1 h at room temperature followed by 3 consecutive washes in PBS containing 0.1% Triton X-100. The coverslips were next mounted in PPDM onto coverslips.

The following primary antibodies were used for immunofluorescence and Western blotting: PLK1 (1:200, SantaCruz: sc-17783), BUB1 (1:200, Abcam: ab54893), anti-centromere antibodies (ACA) (1:200, Antibodies, Inc.: 15-234), INCENP (1:1000, Abcam: ab36453). Microtubules were stained with DM1α (1:1000 IF, 1:10,000 WB, Sigma:T6199). Generation of the CENP-O-P antibody was previously described and was prepared against full length CENP-O/P-His expressed in *Escherichia coli* (McKinley *et al.*, 2015) and used at 1 μg/mL. Generation of the NDC80 “Bonsai” antibody was previously described (Schmidt *et al.*, 2012) and used at 1 μg/mL. DNA was visualized using 10 μg/ml Hoechst (Sigma-Aldrich). Cy2, Cy3-, and Cy5-conjugated secondary antibodies were obtained from Jackson Laboratories and used at 1:300. Immunofluorescence cell images were acquired on a DeltaVision Core deconvolution microscope (Applied Precision) equipped with a CoolSnap HQ2 CCD camera and deconvolved where appropriate. Approximately 35 Z-sections were acquired at 0.2 μm steps using a 100X, 1.4 Numerical Aperture (NA) Olympus U-PlanApo objective or a 60×, 1.42 NA Olympus U-PlanApo objective.

For Western analysis of CENP-O knockout cells, Western blotting was performed on 12% SDS-PAGE gels using 1-hour semi-dry transfer with 3% bovine serum albumin (BSA, Sigma) in TBS + 0.5% Tween-20 as a blocking agent.

### Quantification and Statistical Analysis

Quantification of fluorescence intensity was conducted on unprocessed, maximally projected images using FIJI/image J. For image quantification, all images for comparison were acquired using the same microscope and acquisition settings. For quantification of metaphase alignment, cells were defined as misaligned if at least one off-axis chromosome was observed. Only cells with mature spindle structures were evaluated. Due to the nature of the inducible knockout system, a subset of Cas9 cleavage events will be repaired in a manner that retains the open reading frame resulting in a mixed population of cells. To ensure accurate representation of this mixed population, the first 100 diving cells observed were analyzed from each experimental group, for each biological replicate. For analysis of BUB1, INCENP, and PLK1 intensity at kinetochores, 10 individual kinetochores were selected at random with 4-pixel diameter circles and the total integrated intensity was measured. Background correction was performed by selecting a nearby non-kinetochore region of equal size for each kinetochore and subtracting its integrated intensity from that of the kinetochore region. The average of all kinetochores was then determined per cell, with approximately 20-25 cells analyzed for each condition per experiment. Statistical analyses were performed using Prism (GraphPad Software). Details of statistical tests and sample sizes are provided in the figure legends.

CCAN: Constitutive Centromere Associated Network

## Acknowledgements

We thank the members of the Cheeseman lab, as well as Karen Schindler, for their support and input. This work was supported by grants to IMC from the NIH/National Institute of General Medical Sciences (R35GM126930), National Science Foundation (2029868), and a Pilot award from the Global Consortium for Reproductive Longevity and Equity (GCRLE), and a Damon Runyon post-doctoral fellowship to ALN. The authors declare that they have no conflicts of interest.

## Author Contributions

A.L.N. and I.C. designed the experiments. A.L.N. conducted the experiments. A.L.N. and M.D.F. analyzed the data. A.L.N. and I.C. wrote and edited the manuscript.

## Supplementary figures Legends

**Figure S1.**
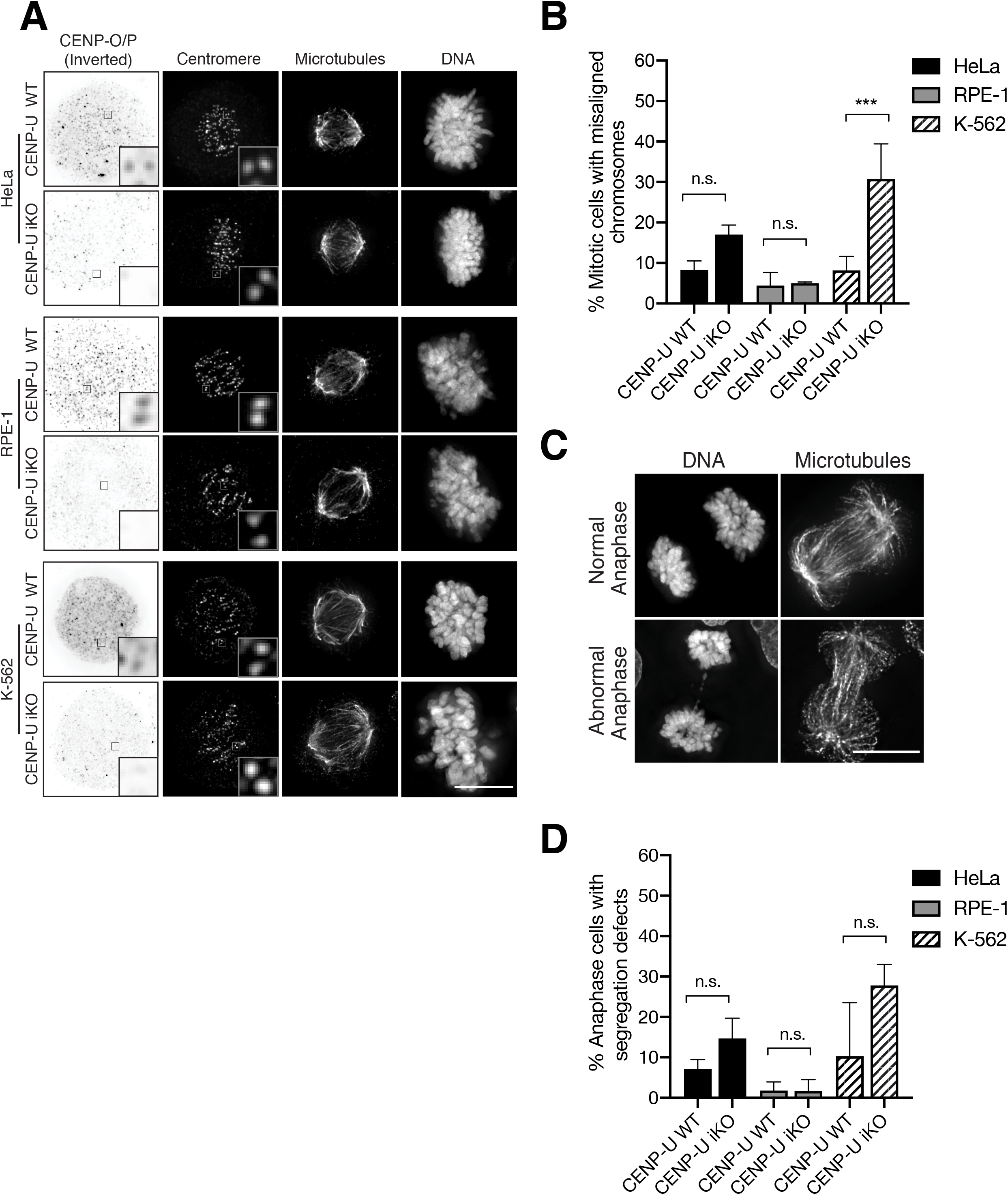
Knockout of CENP-U results in chromosome segregation defects in some but not all cell lines. Related to Figure 1. A) Representative Z-projected immunofluorescence images of metaphase cells from CENP-U inducible knockout HeLa, RPE-1, and K-562 cell lines showing anti-CENP-O/P antibodies (Inverted), centromeres (ACA), microtubules (DM1α), and DNA (Hoechst). Boxes indicate areas of optical zoom.Percent mitotic cells with misaligned chromosomes from A. N = approximately 300 cells per condition across 3 experimental replicates. C) Representative Z-Projected immunofluorescence images of anaphase cells from CENP-U inducible knockout HeLa, RPE-1, and K-562 cell lines showing microtubules (DM1α) and DNA (Hoechst). D) Quantification of anaphase cells with defects including chromosome bridges and lagging chromosomes from C. Representative anaphase cells are from CENP-U control and CENP-U iKO K-562 cell lines. N = approximately 100 cells per condition across 3 experimental replicates. One-way ANOVA was performed (*** = 0.0002). Error bars indicate standard deviation. Scale bars, 10 μM.

**Figure S2.**
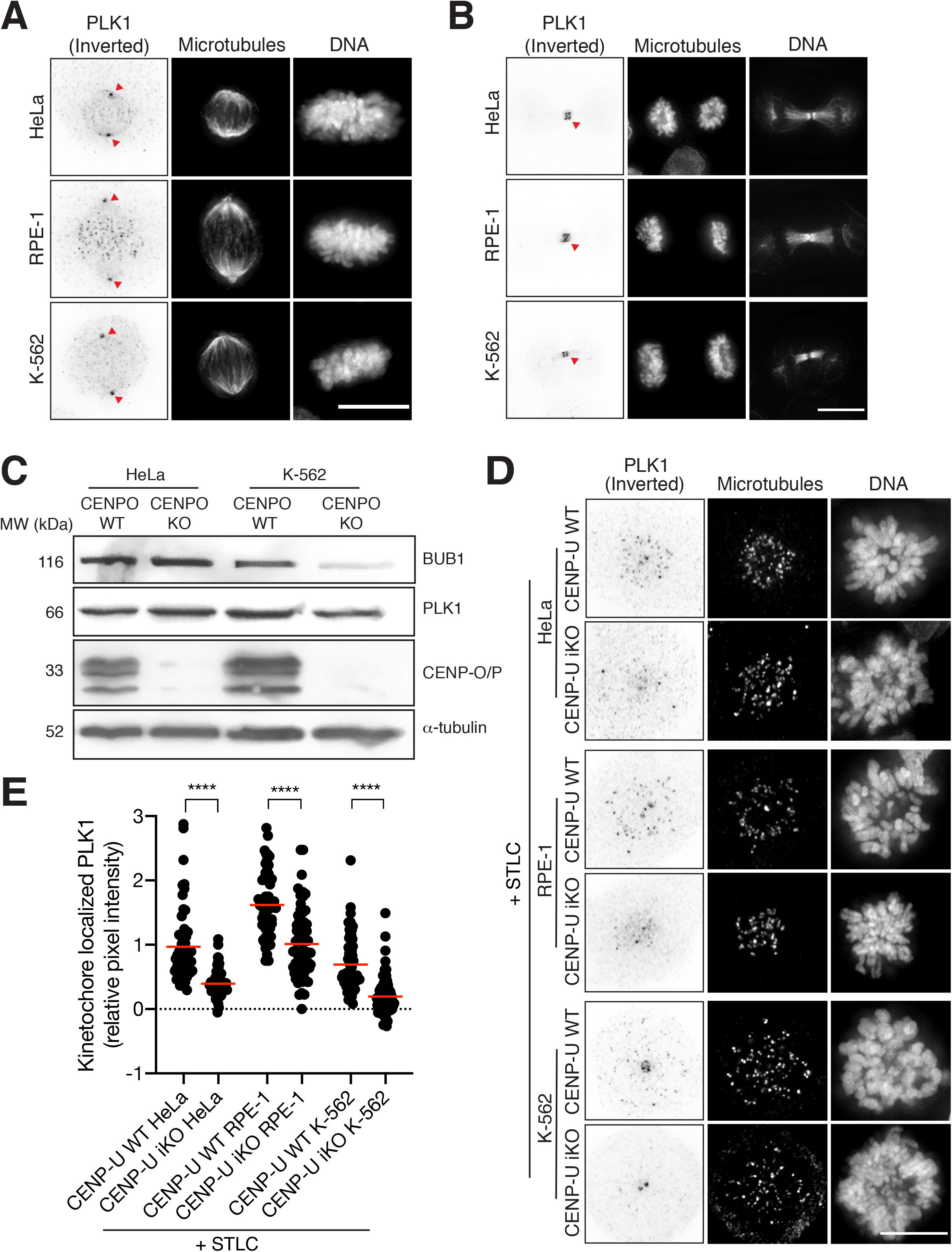
Centrosomal and spindle midzone populations of PLK1 are unperturbed upon CENP-O complex loss. Related to Figure 2. A-B) Representative Z-projected immunofluorescence images of metaphase and anaphase cells from HeLa, RPE-1, and K-562 cell lines respectively showing anti-PLK1 (inverted), centromeres (ACA), and DNA (Hoechst). Arrows indicate centrosomal localized PLK1 (A) and spindle-midzone localized PLK1 (B). C) Western Blots of the indicated proteins from stable clonal CENP-O knockout cell lines from the HeLa and K-562 backgrounds. Clonal cells were isolated after long-term knockout induction and synchronized in mitosis via incubation in the Kif11 inhibitor STLC overnight prior to processing. D) Representative Z-projected immunofluorescence images of STLC-arrested metaphase cells from CENP-U inducible knockout HeLa, RPE-1, and K-562 cell lines showing anti-PLK1 (inverted), centromeres (ACA), and DNA (Hoechst). To ensure a comparison of PLK1 levels at similar stages of mitosis, cells were synchronized via incubation in STLC overnight prior to fixation. E) Relative pixel intensity of kinetochore-localized PLK1, normalized to HeLa control cells, from D. Each data point represents a single cell. Red bars indicate mean. N= Approx. 60 cells per group across 2 experimental replicates. One-way ANOVA was performed (**** = <0.0001). Scale bars, 10 μM.

**Figure S3.**
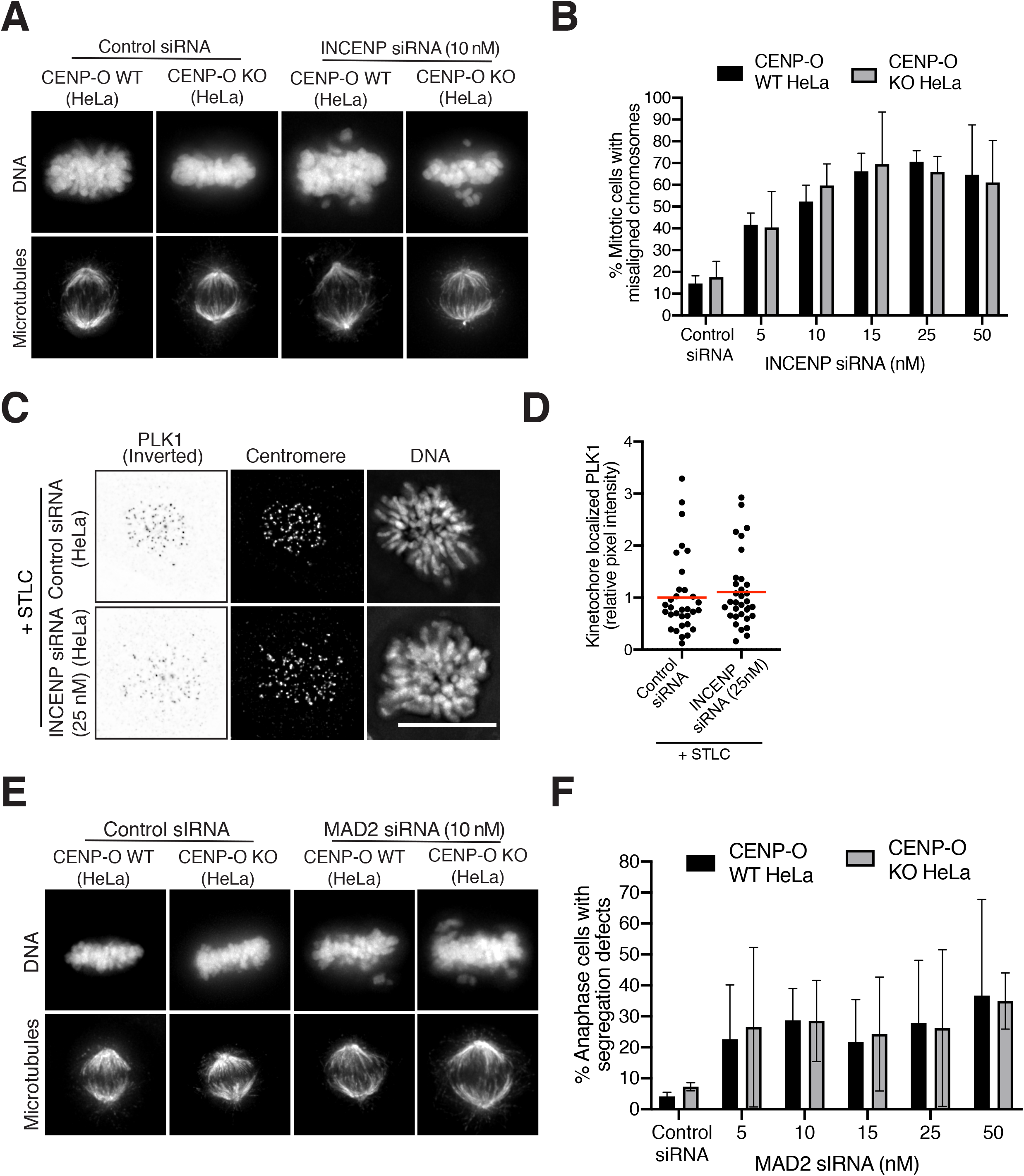
The sensitization of CENP-O knockout cells is specific to BUB1. Related to Figure 3. A-F) HeLa control and stable CENP-O knockout (KO) cells were incubated in the presence the indicated concentrations INCENP or MAD2 siRNAs or non-targeting control siRNAs for 48 hours prior to analysis. A) Z-projected immunofluorescence images of metaphase cells of the indicated cell lines incubated in the presence of control siRNA and 10 nM INCENP siRNA showing microtubules (DM1α) and DNA (Hoechst). B) Percent mitotic cells with misaligned chromosomes from E. N = approximately 300 cells per condition/per group across 3 experimental replicates. Error bars show standard deviation. Two-way ANOVA was performed with no significant difference observed. C)Representative Z-projected immunofluorescence images of STLC-arrested metaphase cells HeLa cells incubated in the presence of control siRNAs or 10 nM INCENP siRNA showing anti-INCENP (inverted), centromeres (ACA), and DNA (Hoechst). To ensure quantification of cells at similar mitotic timepoints, cells were arrested in metaphase via incubation in STLC overnight. D) Relative pixel intensity of kinetochore-localized INCENP, normalized to control siRNA cells, from C. Each data point represents a single cell. Red bars indicate mean. N = 30 cells per group, across 2 experimental replicates. Student’s t-test was performed with no significant difference observed. E) Z-projected immunofluorescence images of metaphase cells of the indicated cell lines incubated in the presence of control siRNA or 10 nM MAD2 siRNAs showing microtubules (DM1α) and DNA (Hoechst). F) Percent mitotic cells with misaligned chromosomes from E. Error bars indicate standard deviation. N = approximately 300 cells per condition/per group across 3 experimental replicates. Two-way ANOVA was performed with not significant difference observed. Scale bars, 10 μM.

## Notes

### Competing Interest Statement

The authors have declared no competing interest.

